# Direct label-free imaging of nanodomains in biomimetic and biological membranes by cryogenic electron microscopy

**DOI:** 10.1101/2020.02.05.935551

**Authors:** Frederick A. Heberle, Milka Doktorova, Haden L. Scott, Allison Skinkle, M. Neal Waxham, Ilya Levental

**Affiliations:** University of Tennessee; University of Texas Health Science Center at Houston; University of North Carolina at Chapel Hill; McGovern Medical School

## Abstract

The nanoscale organization of biological membranes into structurally and compositionally distinct lateral domains is believed to be central to membrane function. The nature of this organization has remained elusive due to a lack of methods to directly probe nanoscopic membrane features. We show here that cryogenic electron microscopy (cryoEM) can be used to directly image coexisting nanoscopic domains in synthetic and bio-derived membranes without extrinsic probes. Analyzing a series of single-component liposomes composed of synthetic lipids of varying lengths, we demonstrate that cryoEM can distinguish bilayer thickness differences as small as 0.5 Å, comparable to the resolution of small-angle scattering methods. Simulated images from computational models reveal that features in cryoEM images result from a complex interplay between the atomic distribution normal to the plane of the bilayer and imaging parameters. Simulations of phase separated bilayers were used to predict two sources of contrast between coexisting ordered and disordered phases within a single liposome, namely differences in membrane thickness and molecular density. We observe both sources of contrast in biomimetic membranes composed of saturated lipids, unsaturated lipids, and cholesterol. When extended to isolated mammalian plasma membranes, these methods reveal similar nanoscale lateral heterogeneities. The methods reported here for direct, probe-free imaging of nanodomains in unperturbed membranes open new avenues for investigation of nanoscopic membrane organization.

**SIGNIFICANCE:** We have used cryoEM to achieve direct, probe-free imaging of lateral domains in biomimetic lipid membranes under native conditions and to characterize differences in their structures. First, measurements of membrane thickness in laterally uniform single-component membranes show that cryoEM is capable of sub-angstrom resolution of interleaflet membrane thickness. All-atom simulations are used to predict the cryo-EM appearance of submicron domains in vesicles with coexisting liquid domains and these are quantitatively validated by direct imaging of phase separated membranes. We then extend this approach to observe nanoscopic domains in isolated cellular membranes, comprising the first direct imaging of nanodomains in biomembranes.

## INTRODUCTION

Membrane physiology is intrinsically dependent on the structural and dynamical properties of lipid bilayers. Lipid collective behavior influences membrane fluidity, lipid packing, bilayer thickness, elastic moduli, and surface charge density, which in turn affect protein interactions, dynamics, and functions (Janmey and Kinnunen 2006, van Meer, Voelker et al. 2008). A striking example is the variation in membrane physical properties in the eukaryotic secretory system: endoplasmic reticulum (ER) membranes are thinner and more fluid than those of the Golgi, which are themselves thinner and more fluid than plasma membranes (PM) (Mitra, Ubarretxena-Belandia et al. 2004). These thickness differences have been implicated in protein sorting through the secretory pathway, with proteins that contain shorter transmembrane regions being preferentially retained in the ER while longer transmembrane domains (TMDs) are sorted to the PM (Bretscher and Munro 1993, Sharpe, Stevens et al. 2010, Lorent, Diaz-Rohrer et al. 2017).

In addition to variations between *different* membranes, the particular lipid compositions of many eukaryotic cells may be poised for variations *within* a given membrane. Preferential interactions between specific lipid classes (e.g., saturated acyl chains, sphingolipids, glycolipids, and sterols) can combine to produce lateral heterogeneities with unique structures, compositions, and putative cellular functions (Sezgin, Levental et al. 2017). This capacity for functional lipid-driven lateral segregation of biomembranes is known as the ‘lipid raft hypothesis’ and has been one of the most widely studied (Levental and Veatch 2016) and controversial (Munro 2003) topics of modern membrane biology.

The physicochemical underpinning of the raft hypothesis is the fact that biomimetic mixtures of synthetic or isolated lipids often form coexisting liquid phases (reviewed in (Feigenson 2007, Marsh 2009)), with the cholesterol- and saturated-lipid-rich liquid-ordered (Lo) phase serving as the model for biological lipid rafts. Phase separated model membranes have yielded a wealth of insights into the principles underlying the organization of biological membranes; however, their lack of proteins and compositional complexity prevents a straightforward translation to living membranes. Within that context, the direct observation of liquid-liquid phase separation in PMs isolated from living cells as Giant Plasma Membrane Vesicles (GPMVs) was a key confirmation of the capacity of biological membranes to separate into coexisting lipid domains (Baumgart, Hammond et al. 2007), which was later confirmed in yeast vacuoles (Toulmay and Prinz 2013, Rayermann, Rayermann et al. 2017). Since then, GPMVs have been widely used to assay the lipid and protein compositions of raft domains, their resulting physical properties, modulation of these properties by external inputs, and the nature of the miscibility phase transition in biomembranes. For example, it was shown that proteins with longer TMDs preferentially sort into the more tightly packed raft phase (Diaz-Rohrer, Levental et al. 2014), implying that this raft phase is also relatively thick. Most importantly, protein partitioning in GPMVs is strongly related to the sub-cellular localization of proteins in live cells, suggesting a functional role for rafts in membrane traffic (Diaz-Rohrer, Levental et al. 2014). Similar connections between GPMVs and cell physiology have been reported for the diffusion of lipids (Kinoshita, Suzuki et al. 2017), nanoscale protein organization in live cells (Stone and Veatch 2015), and the assembly of viral proteins at the PM (Ono 2010, Sengupta, Seo et al. 2019).

Despite these and other significant advances supporting the functional relevance of ordered membrane domains, the nature and function of lipid rafts remain controversial. Studies using imaging mass spectrometry failed to observe cholesterol-rich domains in cells (Frisz, Lou et al. 2013). Similarly, a single-molecule tracking study reported no effects of enriching putatively raft-associated GPI-anchored proteins on other proteins’ localization or diffusion (Sevcsik, Brameshuber et al. 2015). Ultimately, the controversy persists because direct, microscopic detection of domains in mammalian cells has proven elusive, though it should be emphasized that such domains have been directly observed in yeast (Toulmay and Prinz 2013, Rayermann, Rayermann et al. 2017).

Seeing is believing and therefore some of the most compelling studies of liquid-liquid phase separation have involved direct imaging of micron-sized domains in GUVs and GPMVs (Baumgart, Hess et al. 2003, Veatch and Keller 2003, Veatch, Polozov et al. 2004, Hammond, Heberle et al. 2005, Baumgart, Hammond et al. 2007). However, such large, thermodynamically stable domains are generally not observed in mammalian plasma membranes, where active processes and cytoskeletal interactions (Sharma, Varma et al. 2004, Plowman, Muncke et al. 2005) may be expected to produce more dynamic and nanoscopic structures. Thus, tools to investigate the nanoscale organization of biological and biomimetic membranes are essential. Several methods, including ESR (Ionova, Livshits et al. 2012), NMR (Veatch, Polozov et al. 2004, Juhasz, Sharom et al. 2009), SANS (Heberle, Petruzielo et al. 2013), FRET (Heberle, Wu et al. 2010), and interferometric scattering microscopy (de Wit, Danial et al. 2015) have proven useful for indirectly probing nanoscale phase separation across temporal and spatial scales. In principle, both AFM and super-resolution microscopy are capable of directly detecting domains below the optical resolution limit, but both require immobilizing membranes on solid supports, which inherently affects bilayer structure and dynamics (Scomparin, Lecuyer et al. 2009, Goodchild, Walsh et al. 2019). To date, there has been no direct imaging of coexisting nanoscopic phases in unsupported membranes, including synthetic biomimetic models, isolated biomembranes, or cell membranes *in situ*. Consequently, major questions remain about the sizes, morphologies, physical properties, composition, and very existence of membrane nanodomains.

To address some of these questions, we have applied cryoEM, a revolutionary development in structural biology that allows direct, probe-free, angstrom-level resolution of biological samples *in situ* (Egelman 2016)(Fischer, Dash et al. 2018). Previous observations have suggested that EM is sensitive enough to resolve the two bilayer leaflets, with cell membranes appearing as a double line separated by about 3-5 nm in EM images (Fischer, Dash et al. 2018, Lewis, Caldara et al. 2018). What has not been demonstrated is whether cryoEM is capable of resolving the known thickness difference between coexisting Ld and Lo phases in model membranes or analogously hypothesized thickness differences in biomembranes. Here, we show that cryoEM can provide sub-angstrom resolution of membrane thickness differences and that these can be used to directly image nanodomains in biomimetic and biological membranes. These (and the jointly-submitted manuscript by Cornell et al.) are, to our knowledge, the first direct observations of nanodomains in free-standing biomimetic membranes.

## RESULTS

### CryoEM detects bilayer thickness differences with sub-angstrom resolution

To assess the capability of cryoEM to measure bilayer thickness, we directly imaged unstained synthetic lipid membranes. Figure 1A shows a cryoEM image of vesicles composed of dioleoyl phosphatidylcholine (DOPC; di18:1-PC), a phospholipid that spontaneously forms bilayers upon hydration. Extrusion through a 50 nm polycarbonate filter produces largely monodisperse vesicles with an average diameter of 74 ± 10 nm (relative polydispersity = 0.14) as determined from the images (unless noted, all synthetic vesicles experimentally examined in this paper contain 5 mol% of the negatively-charged lipid POPG to minimize multilamellar vesicles (Scott, Skinkle et al. 2019)). A large majority of the extruded vesicles consisted of a single bilayer, the structure of which is evident as two dark concentric rings (referred to here and throughout as troughs) flanked by brighter bands (Fig. 1A, *lower left*). This intensity variation arises from phase contrast between the transmitted and scattered electron waves, as previously described (Wang, Bose et al. 2006).

**Figure 1.**
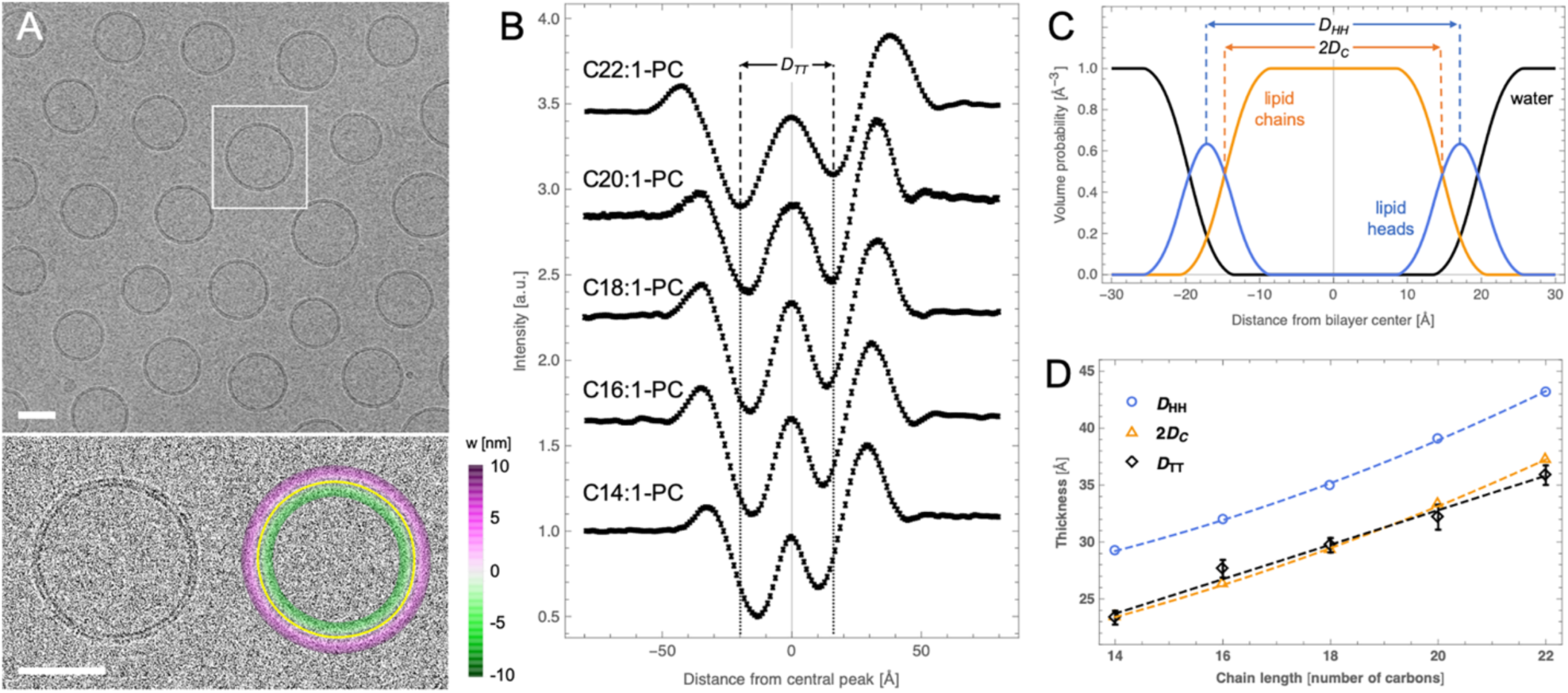
CryoEM is sensitive to sub-angstrom variation in bilayer thickness. (A) CryoEM image of a field of di18:1-PC vesicles prepared at room temperature (*upper*) and an expanded view of a vesicle without (*lower left*) and with (*lower right*) an overlay of the fitted vesicle contour (scale bars 50 nm). The color scale in the lower right image corresponds to the length of the shortest line connecting each pixel to the contour (i.e., its distance from the bilayer center). (B) Average intensity profiles from cryoEM images for a series of fluid-phase PC vesicles. The trough-to-trough distance D_TT_ is shown for the C22:1 bilayer and extended to the *x*-axis to show relative differences between the profiles. (C) The volume probability profile for a di18:1-PC bilayer obtained from analysis of SAXS data at 25 °C. Shown are two standard reference thicknesses defined by SAXS analysis: the headgroup-to-headgroup distance D_HH_, and the hydrocarbon thickness 2D_C_. (D) Bilayer thicknesses determined from SAXS (D_HH_ and 2D_C_) and cryoEM (D_TT_) for the PC series (dashed lines are a visual guide). D_TT_ is shown as average ± SEM of ∼ 50 vesicles. The uncertainty of SAXS thicknesses are estimated to be ± 2%. All vesicles contained 5 mol% POPG to minimize multilamellar structures.

To quantify these features, we designed and implemented an algorithm to determine the intensity profile along the bilayer normal (see Supporting Information for details). Briefly, the vesicle contour is arbitrarily identified as the bright central peak between the dark troughs (Fig. 1A, *lower right*, solid yellow line). Then, individual pixel intensities within 10 nm of the contour are binned by distance from the nearest contour point to generate radially integrated intensity profiles I(w) (Fig. 1B). To test how such profiles reflect differences in membrane thickness, we imaged vesicles comprising a series of PC lipids with different acyl chains ranging from 14 to 22 carbons in length, all containing one unsaturation per acyl chain to ensure fluid phase at room temperature (at which all samples were cryopreserved). Figure 1B shows I(w) profiles (averaged over 5 nm arc-length segments from > 50 vesicles) for this lipid series. The trough-to-trough distance D_TT_ systematically increases with increasing chain length (Table 1, black symbols in Fig. 1D), fully consistent with the expected increase in bilayer thickness.

**Table 1.** Structural parameters derived from analysis of experimental cryo-EM images. Samples are fluid-phase bilayers composed of PC lipids with two monounsaturated chains and cryopreserved at room temperature. D_TT_ corresponds to the measured distance between the troughs of the I(w) profile obtained by averaging the profiles of 5 nm arc-length bilayer segments from > 50 vesicles. Uncertainty in D_TT_ was estimated using standard Monte Carlo methods as described in Supporting Information.

| Lipid | $N_{ves}$ | $N_{seg}$ | $D_{TT}$ [$\text{\AA}$] |
| --- | --- | --- | --- |
| di14:1 | 56 | 2244 | $23.4 \pm 0.6$ |
| di16:1 | 50 | 2211 | $27.7 \pm 0.8$ |
| di18:1 | 98 | 4256 | $29.7 \pm 0.6$ |
| di20:1 | 48 | 1489 | $32.2 \pm 1.1$ |
| di22:1 | 51 | 2820 | $35.9 \pm 0.9$ |

We hypothesized that, as an electron scattering technique, cryoEM should be especially sensitive to the electron-rich phosphates, and that D_TT_ should therefore roughly correspond to the distance between lipid headgroups. To test this hypothesis, we compared cryoEM profiles to different bilayer thicknesses (i.e., corresponding to different reference planes within the lipid interfacial region) derived from small-angle X-ray scattering (SAXS) for the same PC series (Fig. 1D, Table S2). X-rays scatter from the electron cloud and are thus particularly sensitive to the location of the headgroup phosphates, the most electron-dense region of a phospholipid bilayer. Fitting scattering curves (shown in Fig. S1) to a model of the lipid volume probability profile along the bilayer normal (shown in Fig. 1C for DOPC) yields several useful bilayer structural parameters including the headgroup-headgroup distance D_HH_ and the hydrocarbon thickness 2D_C_ (Table S2). Somewhat surprisingly, we observed excellent correspondence between D_TT_ and the lipid hydrophobic thickness 2D_C_, rather than the expected distance between the electron-rich phosphate groups D_HH_. Possible explanations for these discrepancies are discussed below, but these data clearly demonstrate that D_TT_ is strongly correlated with bilayer thickness and that cryoEM can detect variation in bilayer thickness with sub-angstrom sensitivity.

### Trends in cryoEM-derived thickness are recapitulated in simulated images

The lack of agreement between D_TT_ and the well-understood headgroup-headgroup distance D_HH_ derived from scattering measurements prompted further investigation into the relationships between bilayer structure and features observed in cryoEM images. To that end, we used all-atom molecular dynamics simulations to predict I(w) profiles for the same PC bilayer series as experimentally probed above. Although simulated membranes were slightly less densely packed than corresponding experimental bilayers resulting in slightly larger lipid areas and thinner bilayers for all compositions (Table S3), trends in thicknesses with increasing chain length were in excellent agreement with those derived from experimental scattering data (compare Fig. 1D and Fig. 2D). Following the procedure described by Wang et al. (Wang, Bose et al. 2006), cryoEM images were computed from all-atom simulations as described in the Supporting Information. Briefly, the image intensity is proportional to the phase shift experienced by an electron passing through the sample. Flat bilayer electron phase shift profiles (*γ*(*w*), with *w* = 0 corresponding to the bilayer center) were computed from time-averaged atomic number density distributions along the bilayer normal. The flat bilayer profiles were converted to radial electron phase shift profiles *γ*(*R*) by adding an arbitrary vesicle radius (taken to be 60 nm) to the *w*-coordinate. The radial profile was then analytically projected onto a plane and convolved with a contrast transfer function (CTF) to produce a synthetic cryoEM image.

**Figure 2.**
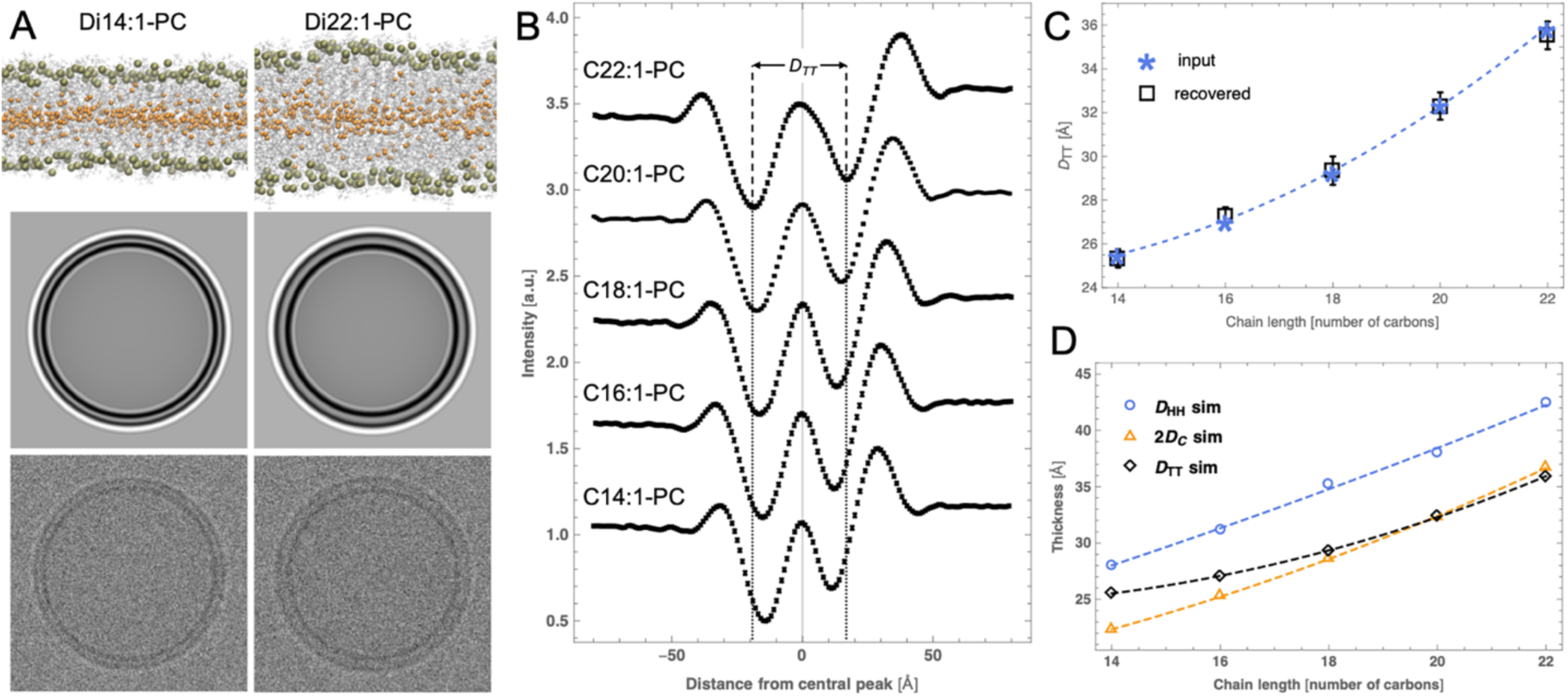
Simulated cryoEM images generated from all-atom bilayer models corresponding to experiments. (A) Snapshots of all-atom molecular dynamics simulations of C14:1-PC and C22:1-PC bilayers (*upper*) with corresponding simulated cryoEM images before (*middle*) and after (*lower*) addition of noise. (B) Average intensity profiles obtained from noisy simulated cryoEM images (N = 60) for a series of PC lipids. The vertical dashed lines show the trough-to-trough distance D_TT_ for the 22:1 bilayer to highlight differences between the profiles. (C) D_TT_ calculated from analytical I(w) profiles used as input to generate the images (stars) are compared to D_TT_ (open squares with error bars) recovered from analysis of noisy simulated images. (D) Comparison of the chain length dependence of bilayer thicknesses (defined in Fig. 1 caption) and D_TT_, all derived from simulations. Dashed lines in panels C and D are visual guides.

Simulated cryoEM images (without and with added Gaussian noise), shown in Fig. 2A for the thinnest (di14:1-PC) and thickest (di22:1-PC) bilayers in the series, faithfully reproduce the pattern of alternating light and dark rings observed in experimental images and reveal a distinctly larger D_TT_ for the thicker bilayer. Average I(w) profiles obtained from analysis of noisy images (N = 60 vesicles) are shown in Fig. 2B (values reported in Table S4) and show very good overall agreement with the experimentally determined profiles (compare Fig. 1B and Fig. 2B), most relevantly a systematic increase in D_TT_ with increasing lipid chain length. Importantly, the custom image processing algorithm, used to analyze both the experimental and simulation image data, yields an unbiased estimate of the true D_TT_ that is accurate to ± 0.5 Å (Fig. 2C, Table S4).

The comparison between D_TT_ calculated from simulated images and the various bilayer thicknesses derived directly from the all-atom simulations is shown in Fig. 2D. Simulated D_TT_ increases monotonically but not linearly with lipid chain length, in contrast to the trend observed in experimental data (Fig. 1D). In particular, simulated D_TT_ for the shorter chain lengths 14:1 and 16:1 is slightly thicker than the hydrophobic thickness 2D_C_, while in experiments D_TT_ and 2D_C_ are nearly identical (Fig. 1D). While possible reasons for the discrepancy are discussed below, these results confirm the origin of the structural features recovered from the analysis of the cryoEM data, making the approach applicable to the visualization and analysis of nanoscale bilayer organization.

### All-atom simulations predict two sources of contrast between coexisting bilayer phases

Having demonstrated the sensitivity of cryoEM to bilayer thickness through a combination of experiment and simulation, we next sought to determine if cryoEM can resolve coexisting phases within an individual vesicle by differences in their thickness. Previous work has established that Lo phases enriched in cholesterol and high-melting saturated lipids are 6-10 Å thicker than Ld phases enriched in low-melting lipids such as DOPC or POPC (Heberle, Petruzielo et al. 2013, Heftberger, Kollmitzer et al. 2015, Bleecker, Cox et al. 2016). In good agreement with experimental data, a thickness difference of ∼ 8 Å was observed in MD simulations of Ld and Lo phases composed of DPPC, DOPC and cholesterol (Sodt, Sandar et al. 2014). We used simulation trajectories of pure Ld and Lo phases obtained from (Sodt, Sandar et al. 2014) to investigate the ability of cryoEM to detect two environments of different thickness within one vesicle (see Supporting Information for details of the simulations). Analytical projection and convolution of electron phase shift profiles for the two simulated pure Ld and Lo bilayers yielded a D_TT_ difference of 2.3 Å (Fig. 3A), substantially smaller than the 6.7 Å hydrophobic thickness difference between the two bilayers calculated directly from the simulation trajectories (Table S5).

**Figure 3.**
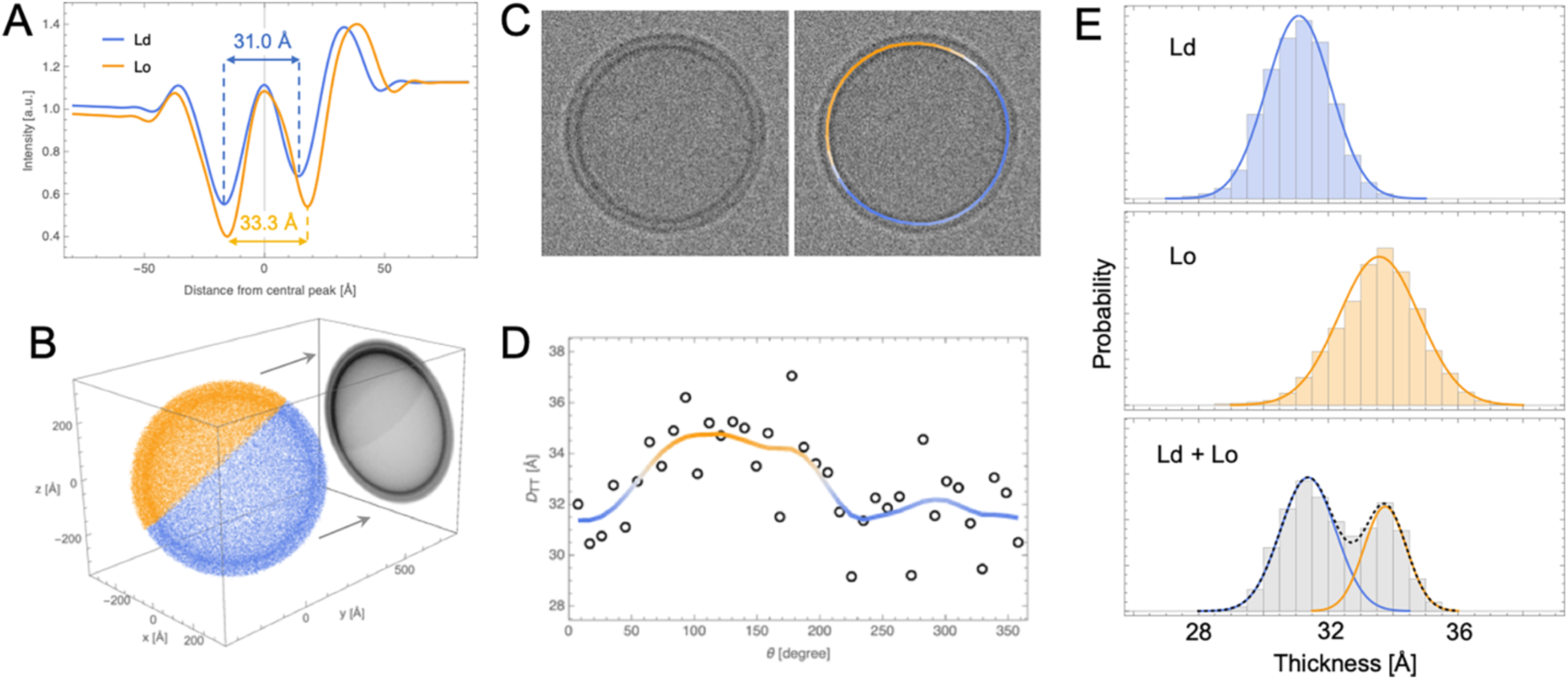
Analysis of simulated cryoEM images of phase separated bilayers. (A) Analytical I(w) profiles calculated from all-atom molecular dynamics simulation trajectories of Lo and Ld phases composed of DPPC/DOPC/cholesterol (Sodt, Sandar et al. 2014). (B) A 2D image of a vesicle with coexisting Ld (blue) and Lo (orange) domains is obtained by projecting a pointillist representation of the 3D spatial variation in electron phase shift density (given by the radial profiles in *A*) onto the *xz* plane. (C) A simulated cryoEM image obtained from the projection in *B* after convolution with a contrast transfer function and addition of noise (*left*) and the same image color-coded by the smoothed thickness values of individual 5 nm arc-length segments around the vesicle contour (*right*). (D) Variation in the thickness of 5 nm segments (open circles) around the circumference of the vesicle in *C*. Local Gaussian smoothing (solid line) reveals the location of Ld and Lo domains. (E) Thickness distribution of 5 nm segments obtained from 100 simulated images of an Ld vesicle (*top*), an Lo vesicle (*center*), and a vesicle with a 60/40 Ld/Lo phase area fraction (*bottom*). Solid lines are fits to Gaussians.

After analyzing the two phases individually, we computationally constructed vesicles that contained coexisting Lo and Ld domains to determine the feasibility of detecting differences between phases within a single vesicle. The analytical procedure described above is not applicable to bilayers that exhibit spatial variation in lipid composition, such as would be expected in a phase-separated vesicle. We therefore constructed images of vesicles with coexisting Ld and Lo phases by modifying a previously described Monte Carlo method for computing scattering curves from such vesicles (Heberle and Anghel 2015) (full details in Supporting Information). Briefly, a 3D pointillist representation of a spherical vesicle with a 60/40 Ld/Lo phase fraction was created by generating random points in proportion to the 3D spatial variation in electron phase shift contrast (Fig. 3B). The set of points representing the phase separated vesicle was then rotated around a randomly oriented vector originating at the center of the vesicle and projected onto the *xz* plane. The projected (2D) points were binned into pixels of 2.5 Å edge length (Fig. 3B), and the corresponding image was created by assigning to each pixel an intensity proportional to the total number of points in the bin. The image was then convolved with a CTF and subjected to Gaussian noise to simulate a cryoEM projection image of a phase separated vesicle (Fig. 3C, left). Figs. 3A-C reveal that differences in the electron phase shift profiles of Ld and Lo bilayers manifest in images as differences in the relative intensities of Ld and Lo troughs (i.e., Lo is darker than Ld) as well as differences in the trough-trough distance (i.e., 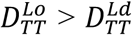) that are visible even in noisy images (Fig. 3C, left).

From the simulated images, we next calculated local D_TT_ for annular segments of ∼ 5 nm arc length around the circumference of each vesicle (Fig. 3D). Although D_TT_ of individual 5 nm segments is somewhat noisy, a clear trend emerges after 4-point Gaussian smoothing (solid line) that is consistent with the location of Ld and Lo domains in the projection (Fig. 3B-C). Fig. 3E shows D_TT_ histograms obtained from 100 simulated images of either an Ld vesicle (top), an Lo vesicle (center), or different random projections of the phase-separated Ld/Lo vesicle shown in Fig. 3B (bottom). Histograms of the simulated Ld or Lo bilayers show unimodal distributions centered at the expected D_TT_ value for each phase, while a clear bimodal distribution is observed in the analysis of phase separated vesicles. By fitting the histogram data to two Gaussians, we recovered a D_TT_ difference (i.e., 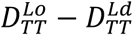) of 2.5 Å (Fig. 3E, bottom), in excellent agreement with the 2.3 Å calculated from an analytical projection of the Ld and Lo phase shift profiles and subsequent convolution with the CTF (Fig. 3A). To summarize, analysis of simulated images suggests that coexisting fluid domains of different thickness within a single nanoscopic vesicle can be readily detected and quantified by cryoEM.

### CryoEM images reveal coexisting nanoscale liquid domains in biomimetic lipid mixtures

A cryoEM image of vesicles composed of the quaternary mixture DPPC/DOPC/POPG/cholesterol 40/35/5/20 is shown in Fig. 4B, with an expanded view of a vesicle shown in Fig. 4D. For comparison, an image of DOPC vesicles (also with 5% POPG) is shown in Fig. 4A, with an expanded view of one vesicle shown in Fig. 4C. Much greater variation in D_TT_ within individual vesicles is observable in the quaternary mixture, consistent with the expected phase behavior of these lipid compositions. Specifically, DOPC forms a uniform fluid Ld phase at room temperature, while ternary mixtures similar to our POPG-doped quaternary mixture separate into coexisting Ld and Lo phases (Veatch and Keller 2003). To our knowledge, these (and images in the jointly-submitted manuscript by Cornell et al.) are the first images of nanodomains in model membranes with coexisting liquid phases under native solvent conditions. D_TT_ histograms obtained from 5 nm bilayer segments of >100 vesicles reveal the expected unimodal distribution for DOPC and a bimodal distribution for the quaternary mixture (Fig. 4E) with a D_TT_ difference (i.e., 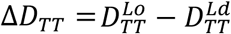) of 7.4 Å and an Ld phase area fraction of 0.64 (Table 2). Ternary mixtures composed of DPPC/DOPC/cholesterol 40/40/20 (i.e., without POPG) also showed a bimodal thickness distribution with Δ*D*_*TT*_ = 6.3 Å and Ld phase area fraction of 0.80 (each calculated as the average of two replicate samples, see Fig. 4E and Table 2), confirming that the contrast was observable without charged lipids. These Δ*D*_*TT*_ values are in good agreement with previously reported thickness differences measured by ensemble-averaged scattering approaches (Heberle, Petruzielo et al. 2013, Heftberger, Kollmitzer et al. 2015) or in supported bilayers by AFM (Bleecker, Cox et al. 2016). Interestingly, Δ*D*_*TT*_ was notably larger in experiments than in simulated bilayers (Fig. 3E), which may be related to approximations used to calculate the electron phase shift profiles from simulation trajectories or to deficiencies in the simulation force fields themselves, as discussed below.

**Table 2.**
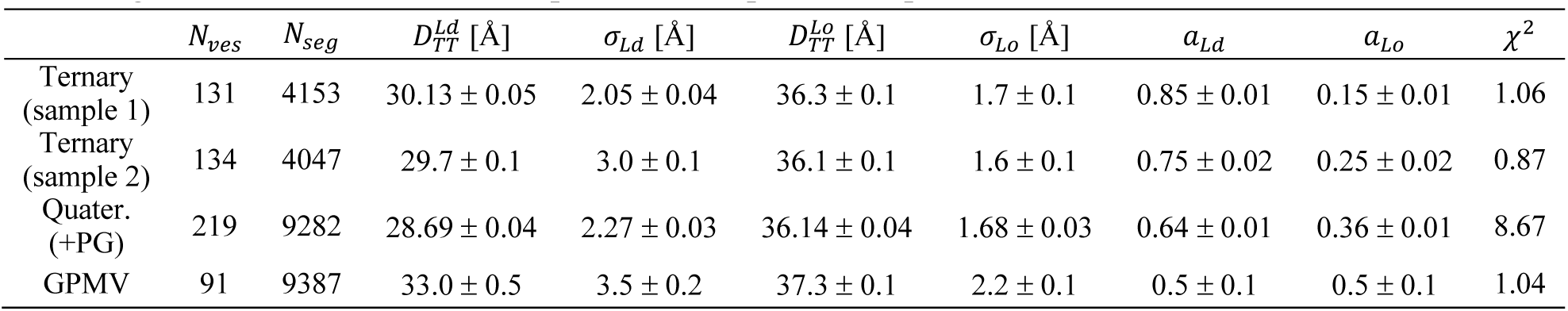
Thickness distribution analysis of multicomponent model membrane vesicles and GPMVs. Parameter uncertainties are calculated from the covariance matrix at the solution of a two-Gaussian fit of the histogram data and do not reflect potential sample-to-sample variation.

**Figure 4.**
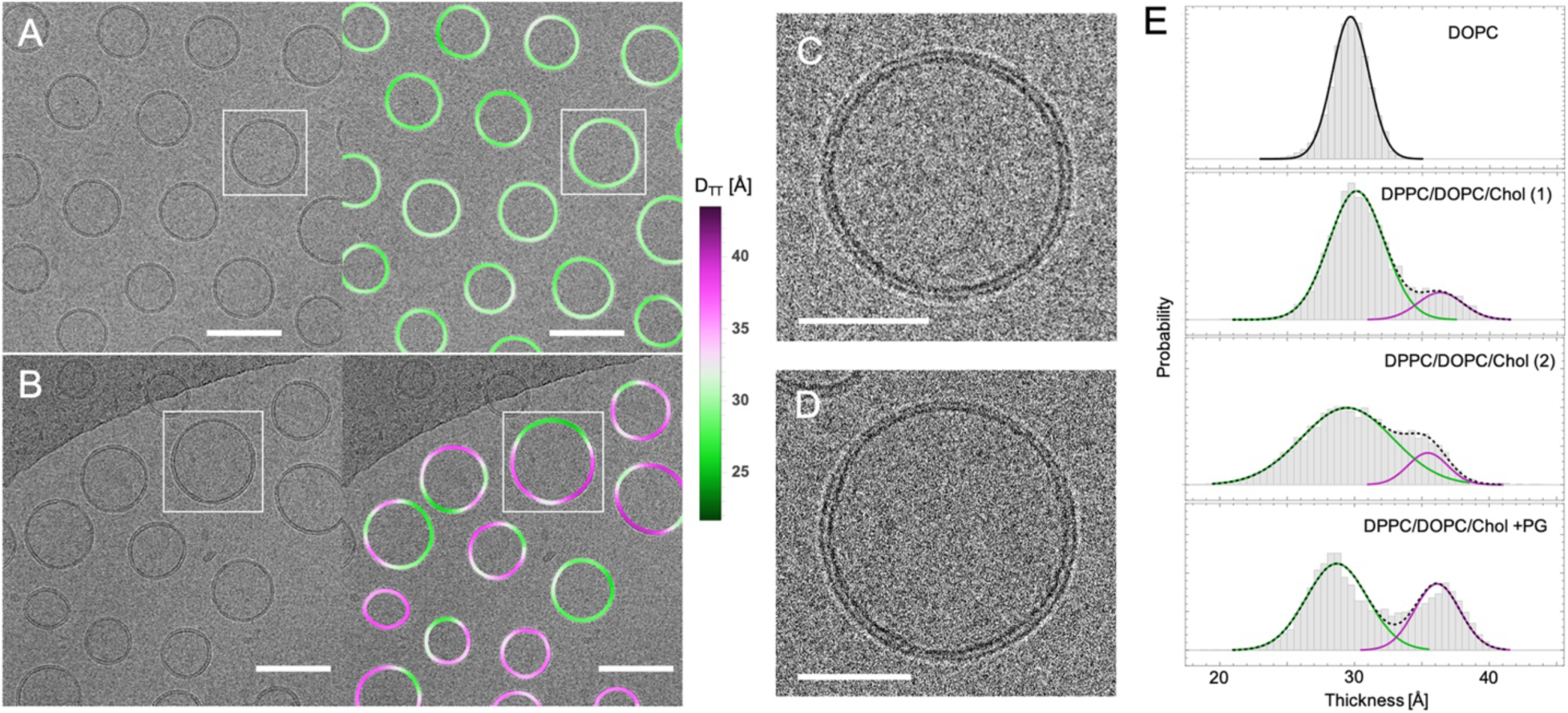
Direct visualization of coexisting phases in cryoEM images of multicomponent lipid mixtures. (A) DOPC vesicles (with 5 mol% POPG) extruded through 50 nm pores (*left*) and the same image color-coded by variation in trough-trough distance D_TT_ (*right*). (B) DPPC/DOPC/POPG/cholesterol 40/35/5/20 vesicles extruded through 50 nm pores (*left*) and the same image color-coded by variation in D_TT_ (*right*), revealing significantly greater thickness variation within individual vesicles as compared to DOPC. Scale bars in panels A and B are 100 nm. (C and D) Zoom-in of boxed vesicles in panels A and B, respectively. Scale bars in panels C and D are 50 nm. (E) Thickness distribution of 5 nm segments obtained from > 100 images of DOPC/POPG 95/5 vesicles (*top*), ternary vesicles with no PG (*middle*), or quaternary vesicles with 5% POPG (*bottom*). Solid lines are fits to Gaussians.

### CryoEM imaging of nanoscale heterogeneity in isolated plasma membrane vesicles

Liquid-liquid phase separation has been directly observed in isolated mammalian PMs (i.e. GPMVs) (Baumgart, Hammond et al. 2007), providing an essential connection between the well-studied phase behavior of simple synthetic membranes and vastly more complex biological membranes (Levental and Levental 2015). While the capacity of isolated plasma membranes to separate into macroscopic Lo/Ld domains has been directly demonstrated, such phase separation is typically observed below room temperature, while at room temperature isolated PM vesicles appear uniform (Baumgart, Hammond et al. 2007, Levental, Lorent et al. 2016, Burns, Wisser et al. 2017, Levental, Surma et al. 2017). How the capacity for phase separation manifests itself under physiological conditions remains unresolved. One experimentally supported hypothesis suggests that PMs are poised for higher-order phase transitions, which produce fluctuations that become smaller (nanoscopic) as temperature increases beyond the macroscopic phase transition (Veatch, Cicuta et al. 2008). To directly evaluate the possibility that isolated PMs retain lateral heterogeneity beyond their macroscopic phase transition temperature, we imaged membrane thickness in GPMVs by cryoEM.

GPMVs were isolated from rat basophilic leukemia (RBL) cells as previously described, concentrated by centrifugation, and cryopreserved at room temperature, similar to the synthetic liposomes above. Although such preparations contain micron-scale vesicles that are typically imaged by light microscopy (Baumgart, Hammond et al. 2007, Sezgin, Kaiser et al. 2012), cryoEM also revealed many 100-500 nm diameter vesicles (Fig. 5). Despite the presence of transmembrane proteins, the bilayers of these vesicles appeared as dark concentric circles with bright halos (Fig. 5A and B), similar in appearance to synthetic liposomes. We applied the above-described methodology to obtain a D_TT_ distribution for 5 nm membrane segments in these isolated PMs that could be compared directly to distributions from synthetic membranes. GPMVs imaged by light microscopy at this temperature appear uniform, as they are above the Lo/Ld miscibility transition temperature (Baumgart, Hammond et al. 2007, Veatch, Cicuta et al. 2008, Levental, Grzybek et al. 2011, Gray, Karslake et al. 2013, Levental, Lorent et al. 2016). However, we observed clearly non-unimodal D_TT_ distributions in GPMV preparations, distinct from all single-component liposome preparations and similar to the phase-separated three-component liposomes (Fig. 5C). The distribution was well described by a two-component fit (Table 2 and Fig. 5C) with relatively thick and thin regions. The thicker areas (37.3 Å) had D_TT_ comparable to the Lo phase, while the relatively thin regions (33.0 Å) were approximately intermediate between the Lo and Ld phase in synthetic membranes. Thus, these images reveal nanoscopic lateral heterogeneities in biomembranes well above the macroscopic phase separation temperature, qualitatively consistent with physical predictions (Veatch, Cicuta et al. 2008).

**Figure 5.**
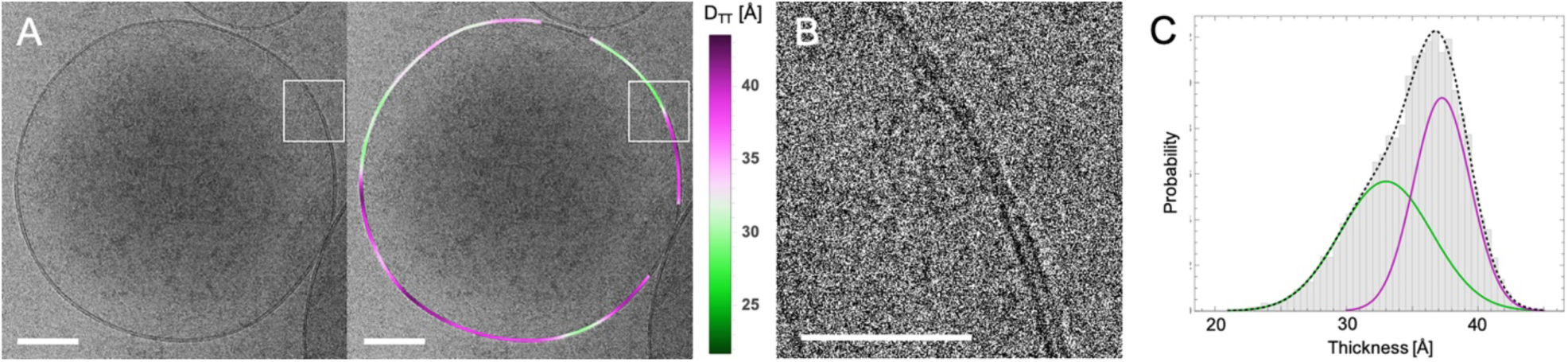
Heterogeneity in GPMVs revealed by cryoEM. (A) GPMV image (*left*) and the same image color-coded by variation in trough-trough distance D_TT_ (*right*). (B) Zoom-in of boxed regions in panel A showing a domain boundary. Scale bar is 100 nm in panel A and 50 nm in panel B. (C) Thickness distribution of 5 nm segments obtained from 91 GPMVs. Solid lines are fits to Gaussians.

## DISCUSSION

### Direct probe-free imaging of nanoscopic domains from thickness differences

We and the jointly-submitted manuscript by Cornell et al. report the first (to our knowledge) direct images of coexisting Ld and Lo nanodomains in unsupported lipid vesicles. No probes were added to the vesicles in this study. Instead, contrast in cryoEM images derives from thickness differences between the coexisting phases. Notably, thickness differences between phases are not essential, as differences in electron density of the two phases also provide intensity contrast that is apparent in images. Very few techniques are capable of resolving liquid-liquid coexistence in free-standing membranes without extrinsic probes. SANS (Heberle, Petruzielo et al. 2013, Marquardt, Heberle et al. 2015) and ^2^H-NMR (Vist and Davis 1990, Veatch, Polozov et al. 2004, Juhasz, Sharom et al. 2009) rely on deuterated lipids, which—while not found in living organisms—are nevertheless true mixture components rather than trace probes. X-ray scattering of oriented bilayer stacks can resolve coexisting Ld and Lo by differences in either the average orientation of chain tilt at wide scattering angles (Mills, Tristram-Nagle et al. 2008) or lamellar repeat distance at small scattering angles (Uppamoochikkal, Tristram-Nagle et al. 2010). While the evidence of coexisting phases provided by these spectroscopic/scattering techniques is convincing, it is also indirect and ensemble averaged. More importantly, information about domain size and properties is either entirely lacking or relies on modeling (Heberle, Petruzielo et al. 2013, Heberle and Anghel 2015). The direct imaging of nanodomains demonstrated here can provide insights into domain morphologies and size distributions that may be important for the functions of membrane subdomains in cells.

We note that the nanoscopic size of domains in our images of synthetic vesicles (Fig. 4B) is not an intrinsic property of the lipid mixture: micron-sized vesicles of the same composition show micron-sized Ld and Lo domains (Veatch and Keller 2003). Instead, domain size in these images is constrained by the nanoscopic dimensions of the extruded vesicles themselves. It remains to be determined whether the sources of contrast we report will also be observed in mixtures with intrinsically nanoscopic domains, such as those near a critical point (Veatch, Soubias et al. 2007) or having high fractions of mixed-chain low-Tm lipid (Konyakhina, Goh et al. 2011, Goh, Amazon et al. 2013). We have previously shown that vesicles containing a mixture of low-Tm lipids such as DOPC and POPC (together with cholesterol and a saturated lipid) can have Lo/Ld thickness mismatches as small as ∼ 6 Å (Heberle, Petruzielo et al. 2013). A correspondingly small difference in D_TT_ of the two phases will certainly diminish, and may perhaps even preclude, the ability to resolve small domains in a noisy cryoEM image. These resolution limits will be the subject of continued investigation; however, observations of thickness heterogeneity in GPMVs that appear uniform by light microscopy (Fig. 5) suggest that intrinsically nanoscopic domains can indeed be observed.

### Heterogeneous thickness distribution in cell-derived PMs

GPMVs derived from the plasma membrane of living cells show a broad thickness distribution that is well approximated by a sum of two Gaussians (Fig. 5). The thicker component has a mean D_TT_ that is remarkably similar to that of the Lo phase in ternary DPPC/DOPC/Chol vesicles (37.3 Å in the GPMV compared to 36.2 Å in the ternary mixture). In contrast, the D_TT_ of the thinner regions (33.0 Å) is notably larger than the Ld phase of the model membranes evaluated here (29.9 Å). These observations are fully consistent with previous reports that the ordered phase in GPMVs has similar properties to synthetic Lo phases, whereas the differences between the coexisting phases is significantly smaller in natural membranes compared to synthetic systems (Kaiser, Lingwood et al. 2009, Levental, Lorent et al. 2016). These differences could be attributable to the transmembrane protein content in GPMVs (Levental, Grzybek et al. 2011) or the high abundance of “hybrid” lipids (one saturated and one unsaturated acyl chain), all of which may attenuate the differences between coexisting ordered and disordered lipid phases compared to synthetic, lipid-only mixtures (Levental and Levental 2015). More importantly, the observation that GPMVs contain nanoscopic lateral domains under conditions at which they appear macroscopically uniform may inform on the mechanisms by which the principles of Lo/Ld phase separation are functionalized by living cells. Detailed comparisons to various existing models (e.g. critical fluctuations) are beyond the scope of the current study but may be accessible with existing methodologies.

### Discrepancies between experimental and simulated images

In both experimental and simulated cryoEM images, D_TT_ increases monotonically with lipid length. However, the trends are markedly different at shorter chain lengths, with simulated values changing less steeply than reference thicknesses derived from SAXS measurements (compare Fig. 1D and Fig. 2D). Thus, while images simulated from molecular dynamics all-atom configurations appear to correctly recapitulate the trend of increasing D_TT_ with increasing bilayer thickness, the sources of contrast in the images may not be completely captured by the simulations or their analysis. This limitation is also evidenced in the analysis of the simulated pure Ld and pure Lo bilayers, where 2D_C_ and D_TT_ are almost identical for the Ld bilayer (31.2 vs 31.0 A) but very different for the Lo bilayer (37.9 vs 33.3 A). One possible source for these discrepancies may be the lack in simulated images of detailed electrostatic interactions that determine electron scattering, namely the use of the shielded Coulomb potential of isolated neutral atoms for calculating the electron phase shift profile. This approximation neglects changes in the electron distribution due to covalent bonds in the lipids. Another potential discrepancy is in the analytical approach for calculating D_TT_ from simulated bilayers. The simulated bilayer patches are flat and therefore their electron phase shift profiles are symmetric about the bilayer midplane. In contrast, vesicles are slightly curved, resulting in subtle differences in lipid packing between the two leaflets that are not present in the simulated images. The packing differences between the two leaflets may be larger for shorter chain lipids, which may explain the subtle discrepancies between D_TT_ and 2D_C_ observed for the 14:1 and 16:1 bilayers. Yet another likely source of error is that the simulation force field used here does not account for electron cloud polarizability. Future developments in simulation methodology, including the use of polarizable force fields, may further improve the quantitative agreement between D_TT_ in simulations and experiments.

### Relationship between bilayer thickness and cryoEM D_*TT*_

Bilayer thickness is a fundamentally important membrane parameter. Thickness is a determinant of membrane permeability (Nagle, Mathai et al. 2008), provides a connection between the membrane’s area compressibility and bending modulus (Rawicz, Olbrich et al. 2000), and defines the relationship between lipid volume and lateral area (Kucerka, Perlmutter et al. 2008). Knowledge of bilayer thickness—defined in symmetric membranes as twice the distance between the bilayer midplane and some reference plane within the lipid molecule, typically in the interfacial region—is crucial for many biophysical models that underlie our understanding of membrane function. Despite this central importance, there are few methods to directly measure bilayer thicknesses, with most relying on ensemble averaging and models to interpret scattering behavior. We show here that cryoEM provides the necessary spatial resolution at the single vesicle level to reliably measure relative bilayer thickness and further, can resolve thickness differences between Lo and Ld phases.

It is also important to point out that while D_TT_ is clearly correlated with bilayer thickness and is sensitive to angstrom-level thickness variation (Fig. 1), it does not appear to directly correspond to a reference plane, such as those determined by X-ray and neutron scattering (Heberle, Pan et al. 2012). Specifically, D_TT_ reported here is derived from an image formed by a 2D projection of the electrostatic potential of a 3D vesicle and subsequent modification of the projected intensity by a contrast transfer function. Each of these processes has the effect of smearing the features related to the matter density distribution. Moreover, the spacing of interference fringes in a cryoEM image, from which D_TT_ is measured, arises from a complex interplay between the underlying bilayer structure and the contrast transfer function, which in turn depends on specific instrument parameters. In other words, changing the experimental settings (most notably, the defocus length) can change D_TT_ and thus the apparent bilayer thickness. Therefore, while trends in D_TT_ are reliable when samples are imaged and analyzed with the same protocols, D_TT_ values as obtained represent a bilayer thickness convolved with the exact parameters of imaging conditions. We are presently pursing a more detailed theoretical and experimental analysis that will ultimately be necessary to derive absolute bilayer structural parameters.

### Limitations of cryoEM for detecting domains

While the methods we describe allow ultrastructural insights into biomimetic and biological membranes, a number of limitations have become evident in the course of these investigations. Unsurprisingly, D_TT_ measurements in small bilayer arcs (as in Figs. 3-5) are quite noisy and accurate estimates of population averages require numerous samples, in analogy with single particle averaging in cryoEM. This approach relies on the assumption of homogeneity, both within and between individual vesicles. The uniformity of any given vesicle population is likely preparation-dependent and rarely well characterized. Especially in multi-component mixtures, significant vesicle-to-vesicle heterogeneities are possible.

While images of single-component vesicles suggest that cryoEM can resolve population thickness differences as small as 0.5 Å, this analysis requires significant averaging. Thus, smaller domains will inherently be more difficult to define and measure. The thickness contrast and domain size limitations of this technique remain to be defined. However, we note that all measurements described here have necessitated relatively small numbers of vesicles, with larger sample sizes likely facilitating higher resolution. On the other hand, there are other sources of noise that are more difficult to control, such as variations in ice thickness or sample degradation due to beam radiation.

## METHODS

### Materials

A list of materials is available in the Supporting Information.

### Preparation of Large Unilamellar Vesicles

Aqueous lipid dispersions at 20 mg/mL were prepared by first mixing appropriate volumes of lipid stocks in organic solvent with a glass Hamilton syringe. The solvent was evaporated with an inert gas stream followed by high vacuum overnight. The dry lipid film was hydrated with ultrapure water above the lipid’s main transition temperature (or for multicomponent mixtures, above the main transition of the highest melting component DPPC) for at least 1 h with intermittent vortex mixing. The resulting multilamellar vesicle (MLV) suspension was subjected to at least 5 freeze/thaw cycles using a −80 °C freezer, then extruded through a polycarbonate filter using a handheld mini-extruder (Avanti Polar Lipids) by passing the suspension through the filter 31 times. Unsaturated lipids were extruded at room temperature, and multicomponent mixtures were extruded at 45 °C. Because different measurement techniques have different optimal concentration ranges, samples were diluted with water or buffer prior to measurement as described in the Supporting Information. Samples for cryoEM measurements were typically cryopreserved and measured within a day of preparation. Samples for small-angle X-ray scattering were measured within a few days of preparation.

### Small-angle X-ray scattering (SAXS)

SAXS measurements were performed using a Rigaku BioSAXS-2000 home source system (Rigaku Americas, The Woodlands, TX) equipped with a HF007 copper rotating anode, a Pilatus 100K 2D detector, and an automatic sample changer. SAXS data were collected at a fixed sample-to-detector distance (SDD) using a silver behenate calibration standard, with a data collection time of 3 h. The one-dimensional scattering intensity *I*(*q*) [*q* = 4*π* sin(*θ*)*/λ*, where *λ* is the X-ray wavelength and 2*θ* is the scattering angle relative to the incident beam] was obtained by radial averaging of the corrected 2D image data, an operation that was performed automatically using Rigaku SAXSLab software. Data were collected in 10-minute frames with each frame processed separately to assess radiation damage; there were no significant changes in the scattering curves over time. Background scattering from water collected at the same temperature was subtracted from each frame, and the background-corrected *I*(*q*) from the individual frames was then averaged, with the standard deviation taken to be the measurement uncertainty and used in weighted least-squares analyses described in the Supporting Information.

### Cryogenic electron microscopy (cryoEM)

To cryopreserve vesicles, 4 μL of a 1 mg/mL sample were applied to a Quantifoil 2/2 carbon coated 200 mesh copper grid that was glow-discharged for 1 min at 15 mA in a Pelco Easi-Glow discharge device. After manual blotting, the grids were plunged from room temperature into liquid ethane cooled with liquid N_2_. Cryo-preserved grids were stored in liquid N_2_ until use. Samples were coded prior to preservation and all sample preparation and image collection was accomplished by an investigator blind to the sample composition. Image collection was accomplished at approximately 2 μm under focus on a FEI Polara G2 operated at 300 kV equipped with a Gatan K2 Summit direct electron detector operated in counting mode. Data collection was performed in a semi-automated fashion using Serial EM software operated in low-dose mode (Mastronarde 2005). Briefly, areas of interest were identified visually and 8×8 montages were collected at low magnification (2400x) at various positions across the grid and then individual areas were marked for automated data collection. Data was collected at 2.7 Å/pixel. Movies of 30 dose-fractionated frames were collected at each target site with the total electron dose being kept to < 20 e^−^/Å^2^. Dose-fractionated movies were drift-corrected with MotionCor2 (Zheng, Palovcak et al. 2017).

### cryoEM data analysis

A detailed description of analysis procedures is found in the Supporting Information. Briefly, the vesicle contour, arbitrarily chosen to be the bright central peak between the dark troughs, was identified from filtered images; from the analysis of simulated images, we conservatively estimate the uncertainty in locating the contour to be ± 2 Å. Individual pixel intensities within 10 nm of the contour were then binned by distance from the nearest contour point to generate a radially integrated intensity profile I(w) for the vesicle, from which the trough-trough distance D_TT_ was determined. For analysis of phase-separated vesicles, the contour was divided into segments of ∼ 5 nm width and D_TT_ determined separately for each segment.

### Molecular Dynamics simulations

All-atom molecular dynamics simulations of five single-component bilayers, phosphatidylcholine with di14:1, di16:1, di18:1, di20:1 or di22:1, were performed with NAMD (Phillips, Braun et al. 2005) using the CHARMM36 force field parameters (Klauda, Venable et al. 2010, Klauda, Monje et al. 2012). All bilayers contained 64 lipids per leaflet, 45 waters per lipid and no ions, and except for C14:1 were constructed and equilibrated with the CHARMM-GUI protocols (Jo, Kim et al. 2008, Jo, Lim et al. 2009, Lee, Cheng et al. 2016). Since C14:1 was not available in CHARMM-GUI at the time of construction, the bilayer was built from a DMPC (C14:0) bilayer as explained in the Supporting Information, then equilibrated with the CHARMM-GUI 5-step protocol by imposing appropriate dihedral angle restraints to preserve the *cis* isomerization of the double bonds. The production runs for all bilayers were performed at constant temperature of 298 K and constant pressure of 1 atm using the same simulation parameters as in (Doktorova, LeVine et al. 2018). Number and electron density profiles of each lipid and water atom in the systems were calculated from the last 680-700 ns of the centered bilayer trajectories with the Density Profile tool in VMD (Humphrey, Dalke et al. 1996).

### GPMV isolation and preparation

Rat basophilic leukemia (RBL) cells were cultured in medium containing 60% Eagle**’**s Minimum Essential Medium (MEM), 30% RPMI, 10% FCS, 100 U/mL penicillin, and 100 μg/mL streptomycin at 37 °C in humidified 5% CO_2_. GPMVs were isolated from RBLs as described (Baumgart, Hammond et al. 2007, Sezgin, Kaiser et al. 2012). Briefly, GPMV formation was induced by 25 mM paraformaldehyde + 2 mM DTT in isotonic GPMV buffer containing NaCl, 10 mM HEPES, and 2mM CaCl_2_, pH 7.4. All GPMVs from one 60-mm cultured dish were combined into a 1.5 mL microcentrifuge tube and concentrated into a loose pellet by centrifugation at 20,000 g for 20 minutes. The pellet was resuspended by gentle pipetting in 20 microliters of GPMV buffer immediately prior to cryopreservation.

## ACKNOWLEDGMENTS

This work was supported by National Science Foundation (NSF) Grant No. MCB-1817929 (to FAH). Funding for IL was provided by the NIH/National Institute of General Medical Sciences (GM114282, GM124072, GM120351), the Volkswagen Foundation (grant 93091), and the Human Frontiers Science Program (RGP0059/2019). Funding for MNW was provided by NIH/National Institute of Neurological Disease and Stroke (R01NS101686) and from the William Wheless III professorship. MD was supported by NIH F32GM134704. SAXS measurements were supported by Department of Energy scientific user facilities at Oak Ridge National Laboratory (ORNL). The computational work used the Extreme Science and Engineering Discovery Environment (XSEDE), account TG-MCB180168. The Polara electron microscope was supported, in part, through the Structural Biology Imaging Center at UTHSC-Houston. The Gatan K2 Summit detector was funded by the NIH (S10OD016279). We thank Alex Sodt and Ed Lyman for providing simulation data. We gratefully acknowledge the members of the Levental lab for extensive discussions and support on this manuscript, as well as the Sarah Keller lab for their critical feedback on our manuscript and collegial/collaborative approach to concurrent publication. Authors have no competing interests.

