## Supplementary Information for "Direct label-free imaging of nanodomains in biomimetic and biological membranes by cryogenic electron microscopy"

The Supporting Material for

**Direct label-free imaging of coexisting nanoscopic fluid phases in biomimetic and biological membranes by cryogenic electron microscopy**

F.A. Heberle, M. Doktorova, H.L. Scott, A. Skinkle, M.N. Waxham, I. Levental

consists of 5 sections, 5 tables, and 6 figures.

### S1. MATERIALS and METHODS

*Materials.* Lipids 1,2-dimyristoleoyl-sn-glycero-3-phosphocholine (di14:1), 1,2- dipalmitoleoyl-sn-glycero-3-phosphocholine (di16:1), 1,2-dioleoyl-sn-glycero-3-phosphocholine (di18:1, DOPC), 1,2-dieicosenoyl-sn-glycero-3-phosphocholine (di20:1), 1,2-dierucoyl-sn-glycero-3-phosphocholine (di22:1), 1,2-dipalmitoyl-sn-glycero-3-phosphocholine (DPPC), 1-palmitoyl-2-oleoyl-sn-glycero-3-phospho-(1'-rac-glycerol), sodium salt (POPG), and cholesterol were purchased from Avanti Polar Lipids (Alabaster, AL) as dry powders and used as supplied. Stock solutions of phospholipids and cholesterol were prepared by weighing powder directly into a volumetric flask and dissolving in HPLC-grade chloroform; two tightly-bound water molecules per phospholipid were assumed present in calculations of molar concentrations. Stocks were stored at −20 °C until use. Ultrapure $H_2O$ was obtained from a High-Q (Wilmette, IL) or Milli-Q Millipore purification system (Burlington, MA).

*Construction of the diC14:1-PC bilayer for MD simulations*. To build an all-atom model of the diC14:1-PC (MYPC) bilayer, we first constructed DMPC (diC14:0-PC) and DYPC (diC16:1-PC) bilayer systems with CHARMM-GUI (Jo 2008, Jo 2009). Each system had 64 lipids per leaflet and 45 water molecules per lipid with no ions. We then removed one hydrogen atom from the C9 and one hydrogen atom from the C10 carbons on each chain of the DMPC lipids to create the MYPC bilayer. Since the double bond in DYPC is located at the same place as in MYPC, between carbons C9 and C10, we mapped each MYPC atom on to the corresponding atom in the DYPC bilayer to ensure that the double bond isomerization in the newly constructed MYPC bilayer was correct. Since DYPC has only two hydrogen atoms attached to carbon C14, we mapped the third terminal hydrogen on each chain of MYPC on to the corresponding C15 atoms in DYPC (the longer CH bond was immediately corrected once the equilibration of the MYPC system started). To ensure that the double bond isomerization in MYPC does not change during the energy minimization steps of the equilibration protocol, we followed CHARMM-GUI's 5-step protocol for equilibration by restraining the dihedral angle defined by carbons C8-C9-C10-C11 on each chain to 0 degrees and gradually releasing the restraints in the course of the 5 equilibration steps.

*Scattering model.* Scattering data were analyzed following (Scott 2019). The experimentally observed scattering intensity from a vesicle suspension can be expressed as the product of a single-bilayer vesicle form factor $P(q)$ and a structure factor $S(q)$ that accounts for density correlations between different bilayers (e.g., the stacked bilayers in a multilamellar vesicle):

$$I(q) = q^{-2} P(q) S(q) \,. \qquad Eq. S1.1$$

For purely unilamellar vesicles such as those used in this study, $S(q)$ = 1. Pencer et al. (Pencer 2006) showed that, for a polydisperse vesicle suspension whose sizes follow a Schulz distribution, $P(q)$ is well-approximated by

$$P(q) \approx P_V(R_m, \sigma, q) |F_B(q)|^2 \,, \qquad Eq. S1.2$$

where $R_m$ is the average vesicle radius, $\sigma$ is the relative size polydispersity, and $P_V(R_m, \sigma, q)$ accounts for the influence of vesicle size and shape on the scattering intensity. $P_V$ was set to unity in our analyses because vesicle size does not influence SAXS data within the experimental $q$ range of this study.

The flat bilayer scattering amplitude $F_B(q)$ in Eq. S1.2 accounts for density correlations within a single bilayer, normal to the plane of the bilayer (i.e., the lipid bilayer structure). We model $F_B$ using a symmetric six-slab volume probability distribution with separate components for the lipid headgroups, combined CH and $CH_2$ groups of the hydrocarbon chains, and terminal $CH_3$ groups at the bilayer midplane (Scott 2019):

$$\begin{aligned} F_B(q) = \frac{2e^{-\frac{(q\sigma_S)^2}{2}}}{qD_H A_L V_T V_W (V_C - 2V_T)} \Big| & V_T\{b_W(A_L D_H - V_H)(V_C - 2V_T) + V_W b_H (V_C - 2V_T) \\ & - V_W A_L D_H (b_C - 2b_T)\} \sin\left(\frac{qV_C}{A_L}\right) \\ & + V_T(V_C - 2V_T)(b_W V_H - b_H V_W) \sin\left(qD_H + \frac{qV_C}{A_L}\right) \\ & + V_W A_L D_H (b_C V_T - b_T V_C) \sin\left(\frac{2qV_T}{A_L}\right) \Big| \,. \qquad Eq.S1.3 \end{aligned}$$

In Eq. S1.3, $V_C$, $V_H$, $V_T$, and $V_W$ are the molecular volumes of the lipid hydrocarbon chains, headgroup, terminal methyl, and water, respectively, with corresponding scattering factors $b_C$, $b_H$, $b_T$, and $b_W$ (Table S1). To mimic the smoothing effects of thermal disorder, the step-like volume probability profile was convoluted with a Gaussian of width $\sigma_S$ = 2.9 Å. Explicit bilayer structural parameters include the area per lipid $A_L$ and the headgroup thickness $D_H$; additional structural parameters are derived from fitted model parameters, the most important of which are the total (Luzzati) bilayer thickness $D_B = 2(V_C + V_H)/A_L$, the hydrocarbon thickness $2D_C = 2V_C/A_L$, and the headgroup-headgroup distance $D_{HH}$ calculated as the distance between headgroup peaks in the volume probability profile. We constrained $V_T$ (53 Å$^3$) to improve the robustness of the fitting routine.

*Scattering analysis.* Scattering data were fit using the model described in the previous section, implemented in a nonlinear least-squares routine using custom code written in Mathematica v11.3.0 (Wolfram Research, Champaign, IL). The adjustable parameters were $A_L$, $V_C$, $D_H$, and a constant background and arbitrary scale factor. These parameters were optimized using a Levenberg-Marquardt algorithm included in Mathematica NonlinearModelFit function.

*CryoEM data analysis.* The cryoEM data analysis is described in detail in sections S2-S5 below.

### S2. Calculating the phase shift profile from an all-atom simulation

The primary mechanism for contrast in a cryoEM image is the phase shift in the electron wavefunction as it passes through the sample, in this case a hydrated lipid bilayer vesicle. This phase shift is proportional to the projected electrostatic potential of the vesicle in the direction of the electron beam. In the weak phase object approximation for defocused imaging, the intensity of the recorded image should vary in proportion to the phase shift, which can be estimated from an atomistic simulation of a flat lipid bilayer. Following Wang et al. (Wang 2006) we consider two contributions to the electron phase shift profile $\gamma(w)$ in the direction normal to the plane of the bilayer. These contributions are the neutral atom phase shift $\gamma_n(w)$ and the electrostatic phase shift $\gamma_e(w)$, where

$$\gamma(w) = \gamma_n(w) + \gamma_e(w). \qquad Eq. S2.1$$

The neutral atom phase shift $\gamma_n$ is calculated from the atomic number density profile of the simulation $\rho_i(w)$ (computed directly from the simulation trajectory as described in the Material and Methods) as

$$\gamma_n(w) = \sigma_e \sum_i V_i \rho_i(w), \qquad Eq. S2.2$$

where $V_i$ is the spatially integrated, shielded coulomb potential for an isolated neutral atom, and is equal to 25, 130, 108, 97, and 267 V Å$^{-3}$ for H, C, N, O and P, respectively (Fig. S2.1B). The parameter $\sigma_e$ accounts for the dependence of the electron phase on the projected potential and is equal to 0.73 mrad V$^{-1}$Å$^{-1}$ for 200 keV electrons (Kirkland 1998). As $\rho_i$ has units of Å$^{-3}$, the phase shift has units of mrad Å$^{-1}$.

The electrostatic phase shift $\gamma_e$ is calculated from electrostatic potential profile $\phi(w)$ as

$$\gamma_e(w) = \sigma_e \phi(w). \qquad Eq. S2.3$$

Figure S2.1A shows the neutral atom and total phase shift profiles.

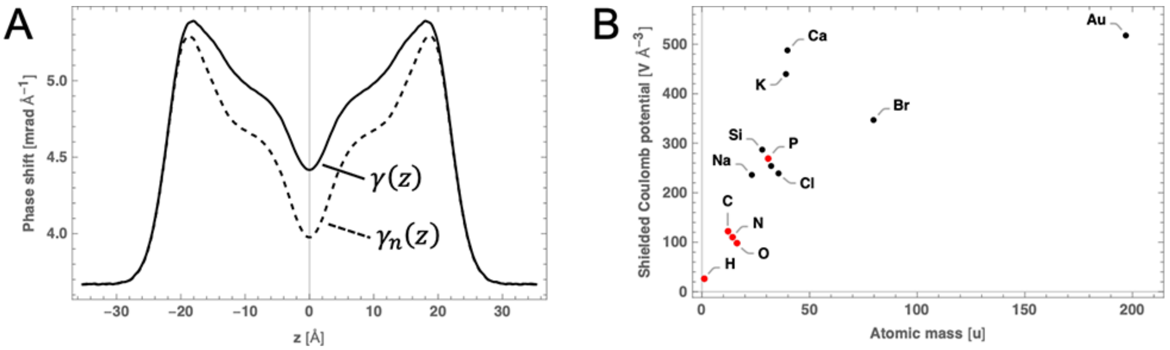


**Figure S2.1 Electron phase shift profiles obtained from molecular simulations.** (A) The total phase shift profile (solid black line) calculated from an all-atom simulation of a DOPC bilayer. The neutral atom component is shown as a dashed black line. (B) The shielded Coulomb potential of isolated neutral atoms.

### S3. Calculating a cryoEM projection image of a vesicle from an all-atom simulation of a uniform lipid bilayer

We now seek to generate a projection image starting from a phase shift profile $\gamma(w)$ calculated in the previous section for a flat simulated bilayer, where $w = 0$ corresponds to the bilayer center. We assume a three-dimensional, spherically symmetric vesicle of radius $R$ with a radial phase shift profile equal to

$$\gamma(r) = \gamma(w + R). \qquad Eq. S3.1$$

A cryoEM image corresponds to a projection of the 3D phase shift variation $\gamma(r)$ onto a plane, followed by convolution with a contrast transfer function (CTF). The projected phase shift profile $\Gamma(r)$ is accomplished by numerical integration of the equation

$$\Gamma(r) = 2\int_r^\infty \frac{\gamma(a - R)}{\sqrt{1 - (r/a)^2}} da. \qquad Eq. S3.2$$

This projected profile is then convolved with a contrast transfer function $c(\boldsymbol{s})$, analogous to the effect of a point spread function in ordinary light microscopy:

$$c(\boldsymbol{s}) = [\sin\chi(\boldsymbol{s}) - Q\cos\chi(\boldsymbol{s})]\exp(-B|\boldsymbol{s}|^2). \qquad Eq. S3.3$$

In Eq. S3.3, $\boldsymbol{s}$ is the spatial frequency in units of Å$^{-1}$, $Q$ is the unitless amplitude contrast factor (typically 5-10%), and $B$ is the amplitude decay factor, typically in the range of hundreds of Å$^2$. The phase perturbation factor $\chi(\boldsymbol{s})$ is given by

$$\chi(\boldsymbol{s}) \approx 2\pi\lambda|\boldsymbol{s}|^2\left(-\frac{\Delta Z}{2} + \frac{C_s\lambda^2|\boldsymbol{s}|^2}{4}\right), \qquad Eq. S3.4$$

where $\lambda$ is the electron wavelength (0.0197 Å for the 300 keV electrons used in this study), $\Delta Z$ is the defocus length (2 μm underfocus in this study), $C_s$ is the spherical aberration coefficient (2 mm for the instrument used here). Neglecting the spherical aberration (which we verified has no discernible effect in our setup) results in the simpler expression

$$\chi(\boldsymbol{s}) \approx -\pi\lambda\Delta Z|\boldsymbol{s}|^2. \qquad Eq. S3.5$$

The reciprocal space image is then given by

$$I(\boldsymbol{s}) \propto 1 + mc(\boldsymbol{s})\mathcal{F}_s[\Gamma(r)](\boldsymbol{s}), \qquad Eq. S3.6$$

where $\mathcal{F}_s$ is the Fourier transform of the projected phase shift profile $\Gamma(r)$ and $m$ is an arbitrary scale factor. The corresponding real-space image is calculated as the inverse Fourier transform of $I(\boldsymbol{s})$,

$$I(\boldsymbol{r}) = \mathcal{F}_r^{-1}[I(\boldsymbol{s})](\boldsymbol{r}). \qquad Eq. S3.7$$

The steps described above are shown schematically in Fig. S3.1

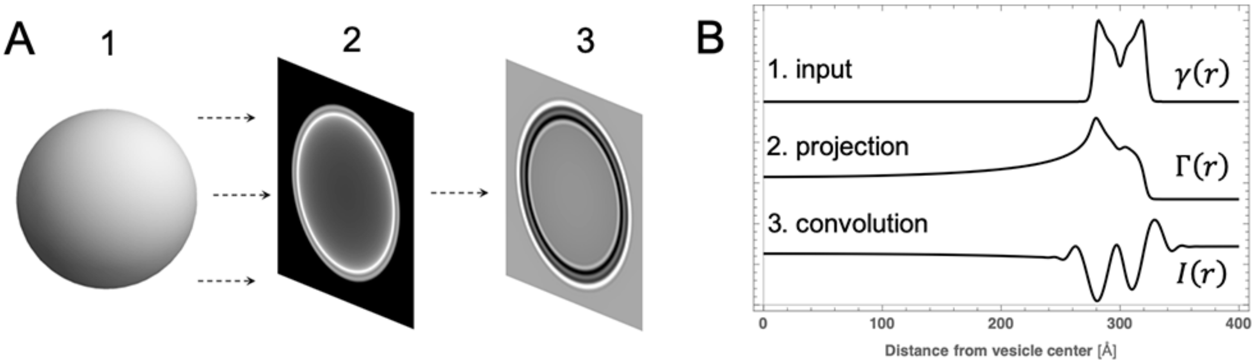


**Figure S3.1 Procedure for generating a cryoEM image from all-atom simulations.** (A) The 3D variation in electron phase shift contrast of a spherical shell (representing a lipid bilayer vesicle) is analytically projected onto a 2D plane (1→ 2) and then convolved with a contrast transfer function (2→ 3). (B) Radially integrated profiles obtained from an input electron phase shift profile (1) after projection (2) and smearing with a CTF (3).

Noise was added to simulated images to better approximate experimentally obtained images. First, the standard deviation $\sigma$ of pixel intensities in experimental images was calculated; a typical value of the relative noise $\sigma/\mu$ was 0.13. Noise was then introduced into the simulated image by sampling individual pixel intensities from a Gaussian distribution centered at the simulated pixel value and with a standard deviation $\sigma$. Examples of noised simulated images are given in Fig. 3A of the main text.

### S4. Simulating an image of a phase-separated vesicle

In the previous section, a simulated phase shift profile of a laterally *uniform* bilayer was analytically projected using Eq. S3.2 to produce $\Gamma(r)$, whose Fourier transform was convolved with a contrast transfer function to produce an image. In the case of a vesicle with coexisting phases (i.e., one that is laterally heterogeneous), we are not aware of a closed form analytical projection analogous to Eq. 3.2. Instead, we modify a Monte Carlo technique originally outlined by Henderson (Henderson 1996) that has successfully been used to compute pair-distance distribution functions for predicting scattering data (Heberle 2015). The method produces a pointillist representation of the vesicle in which the spatial variation in the density of randomly sampled points is proportional to the spatial variation in some physically meaningful bilayer density. In the case of small-angle scattering, it is the electron density contrast (for X-ray scattering) or neutron scattering length density contrast (for neutron scattering) between the lipid bilayer and bulk solvent.

For electron microscopy, the relevant density is the electron phase shift contrast between the bilayer and bulk solvent. In a phase-separated vesicle, the 3D spatial variation in the density $\rho(r, \theta, \phi)$ is completely specified by; (1) the electron phase shift profiles of the Ld and Lo phases (which account for radial variation, i.e. in the direction normal to the bilayer, in each phase); and (2) the location, shape, and size of phase domains (which accounts for the angular variation). We choose as our geometric model a spherical shell centered at the origin with a single round domain centered at $(x, y, z) = (0,0, R)$, where $R$ is the vesicle radius. The domain occupies an area fraction of 0.4. We then perform the following computational steps (which have been generalized for an arbitrary number of domains):

1. The spherical shell representing the vesicle is divided into $j$ subvolumes (not necessarily continuous) of different electron phase shift contrast. For example, a phase-separated vesicle can be represented as a minority phase comprised of one or more small circular domains within a majority matrix phase; the corresponding phase volumes are (1) a set of annular spherical caps (the domains of the minority phase) and (2) a perforated spherical shell (the continuous majority phase). These phase volumes are then further divided into radial shells in which the electron phase shift contrast $\Delta\gamma(r) = \gamma(r) - \gamma_s$, where $\gamma_s$ is the electron phase shift of the solvent (i.e., water) is given by the corresponding electron phase shift profiles of the simulated Ld and Lo phases $\gamma_{Ld}(r)$ and $\gamma_{Lo}(r)$.
2. The volume $V_{i,j}$ and contrast $\Delta\gamma_{i,j}$ of each subvolume is calculated.
3. Within each subvolume, a set of $k$ random, uniformly sampled, three-dimensional coordinates $\boldsymbol{p_{i,j}} = \{(x,y,z)_1, \dots, (x,y,z)_k\}$ is generated, where $k$ is proportional to $V_{i,j}\Delta\gamma_{i,j}$. The constant of proportionality is arbitrary, though it must be the same for all subvolumes, and affects both accuracy and performance (the computational time increases as the square of the number of sampled points).
4. The random points $\boldsymbol{p_{i,j}}$ for all subvolumes are combined in the set $\boldsymbol{p}$ that can be considered a pointillist representation of the electron phase shift contrast in the phase-separated vesicle.
5. The vesicle's orientation is randomized by first generating a random vector $v$ and then rotating each point in $\boldsymbol{p}$ about the origin using the rotation matrix that transforms the vector $u = (0,0,1)$ into $v$.
6. The rotated points are then projected onto an arbitrary 2D plane. A convenient choice is any of the three coordinate planes; for example, projection onto the $xz$ plane corresponds to dropping the $y$-coordinate from each point, i.e. $(x,y,z)_i \to (x,z)_i$.
7. The projected points are binned into pixels of a desired edge length and summed to produce an image $\Gamma(\boldsymbol{r})$.
8. The remaining steps are as given in the previous section (convolution with CTF and addition of noise).

The above procedure is then repeated to produce a set of images that samples the ensemble of projections for randomly oriented vesicles. The next section describes the method for calculating membrane thickness around the projected circumference of an experimental or simulated vesicle.

### S5. Determining the trough-trough distance in cryoEM images

In this section we describe how $D_{TT}$ is determined using simulated images as an example. The first step is to determine the vesicle contour. We found that standard edge detecting algorithms failed in our hands (presumably due to the inherently large noise levels of cryoEM images) which precluded a completely automated fitting routine. Alternatively, by combining local pixel smoothing with an initial guess for the contour, the robustness of the fitting algorithm was dramatically improved as judged by an analysis of simulated noisy images as described below.

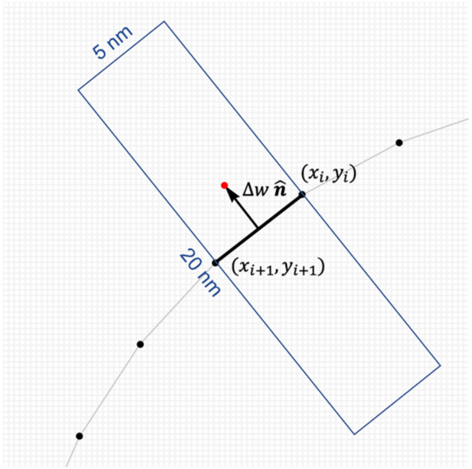


**Figure S5.1 Schematic of the contour fitting algorithm.** Manually drawn contour points, resampled at distances of ~ 5 nm arc length, are shown as black dots. One face of the polygon defined by the manual points is shown as a solid black line. Intensities of pixels contained in a rectangular region extending 10 nm from either side of the face are binned in the direction $\hat{\boldsymbol{n}}$ normal to the face to generate a local intensity profile $I(w)$, where $w = 0$ corresponds to the position of the face. A peak detection algorithm is then used to find the peak closest to the manually drawn face, and its position $\Delta w$ is used to generate the best-fit contour point (shown in red). For comparing distances, a 2.5 Å pixel grid is shown in light gray.

The contour fitting algorithm is shown schematically in Fig. S5.1. For each vesicle, a contour is manually traced on the raw (unfiltered) image using Mathematica's Coordinates Tool. Next, the manually drawn coordinates are smoothed by applying a 3-point Gaussian filter with periodic boundary conditions separately to the $x$- and $y$-coordinates, and the density of points is reduced by resampling at arc length intervals of ~ 5 nm. The resulting set of points constitutes a polygonal representation of the contour. After smoothing the image with a Gaussian filter (5 pixel smoothing radius), all pixels within a 5 × 20 nm rectangular region of interest centered at the polygon face defined by neighboring points $((x_i, y_i), (x_{i+1}, y_{i+1}))$ are selected, and their intensities are binned at 1 Å intervals in the direction normal to the face. The resulting line average represents a local $I(w)$ profile that is approximately normal the bilayer, with $w = 0$ corresponding to the position of the face. Peaks in $I(w)$ are then detected using Mathematica's built-in function FindPeaks. Finally, the best-fit contour point is calculated as

$$\left(\frac{x_i + x_{i+1}}{2}, \frac{y_i + y_{i+1}}{2}\right) + \Delta w\,\hat{\boldsymbol{n}}, \qquad Eq. S5.1$$

where $\Delta w$ is the $w$-coordinate of the peak in $I(w)$ that is closest to $w = 0$, and $\hat{\boldsymbol{n}}$ is the unit vector normal to the face.

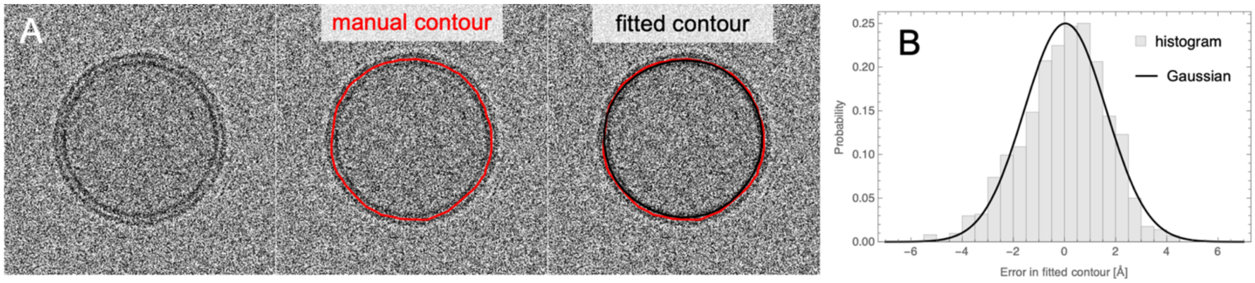


**Figure S5.2 Schematic of the contour fitting algorithm.** (A) Image of a simulated vesicle (*left*) with a manually drawn contour (red line, *center* and *right* images) and a fitted contour (black line, *right* image). (B) Histogram of the error in fitted contour points (calculated as the distance between a best-fit and exact contour point) of 2280 contour points determined from 60 simulated vesicle images. The black line is a fit to a Gaussian.

An example of the contour fitting algorithm described above is shown in Fig. S5.2; the initial manually drawn contour is shown in red, and the final fitted contour is shown in black (Fig. S2A). Simulated images provide a convenient way of estimating the uncertainty in determining the contour, since each simulated projected vesicle is perfectly circular (i.e., the exact contour is known). Figure S2B shows a histogram of the error (calculated as the distance between a best-fit contour point and the corresponding exact contour point) of 2280 contour points determined from 60 simulated vesicle images. The histogram data are well-fit by a Gaussian distribution with a standard deviation of 1.6 Å, which we take to be the approximate uncertainty in determining a contour point from a noisy image.

Final $I(w)$ profiles are determined in a similar procedure, but using the best-fit contour and the unfiltered vesicle image. First, each pixel in the unfiltered image is assigned a $w$-coordinate by calculating its distance to the closest polygon face defined by the set of best-fit contour points. Pixel intensities are then binned by $w$ to generate an average $I(w)$ profile for the vesicle. An example $I(w)$ profile calculated from an image of a simulated phase-separated vesicle is shown in Fig. S5.3 (upper left panel).

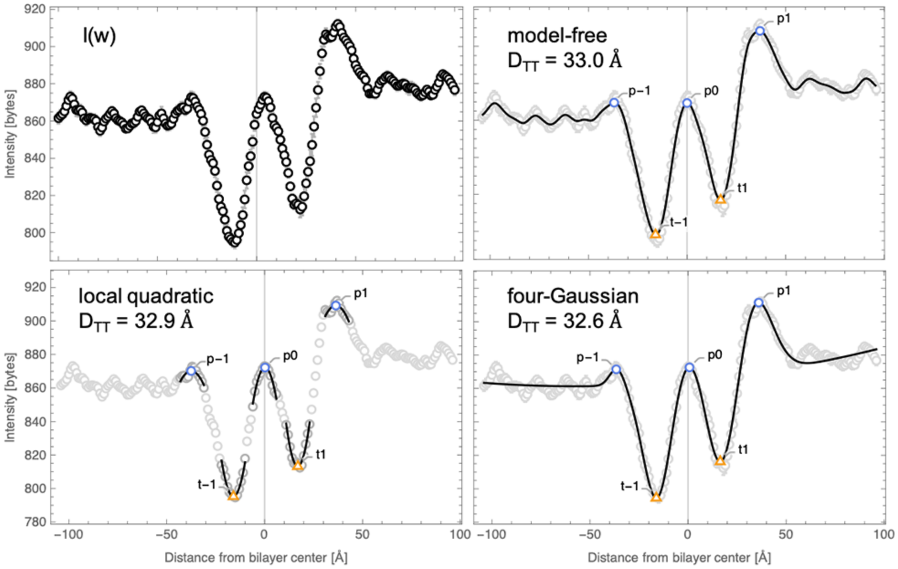

**Figure S5.3 Modeling the intensity profile of an individual vesicle.** *Upper left*, the averaged intensity profile of a single simulated phase separated vesicle. *Upper right*, model-free fit of the profile where the solid line is the raw data Gaussian smoothed with 5 points. *Lower left*, local quadratic fit to the peaks and troughs identified in the model-free fit. *Lower right*, fit of the entire profile to a model consisting of a quadratic background and four Gaussians.

The trough-trough distance $D_{TT}$ is defined as the distance between the $w$-coordinates of the troughs in $I(w)$. We use three methods for locating the troughs. The "model-free" method first performs a local 5-point Gaussian smoothing, and then takes the two absolute minimum intensity values on either side of $w = 0$ to be the troughs (Fig. S5.1, upper right panel). A second "local quadratic" method fits data in the local vicinity of the model-free troughs to quadratic functions and takes the minima to be the troughs. A third "four-Gaussian" method fits the entire profile as a sum of four Gaussians on top of a quadratic background; the troughs then correspond to the two absolute minimum intensity values on either side of $w = 0$. $D_{TT}$ values obtained from the three methods generally agree to within 1 Å, and we report their average.

### SUPPORTING FIGURE

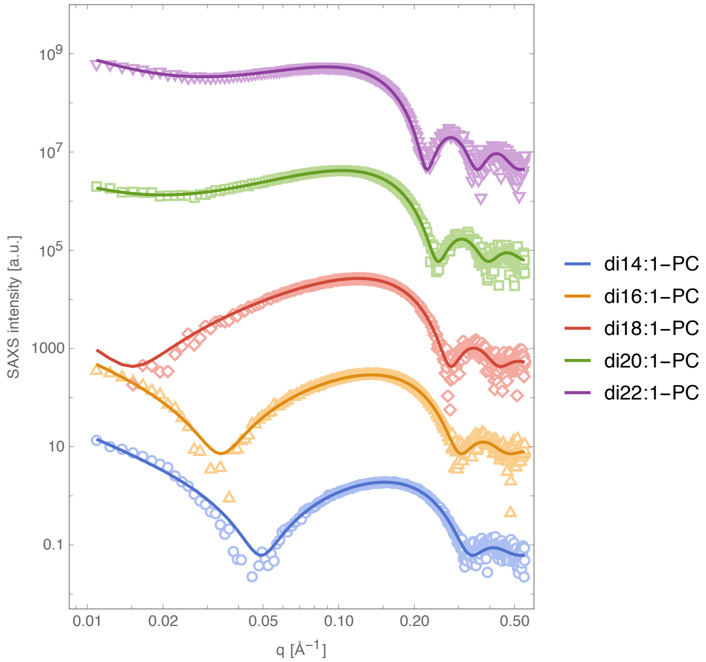


**Figure S1 SAXS data of dimonounsaturated lipid vesicles at 25°C.** Experimental scattering data (open symbols) and fitted curves (solid lines).

### SUPPORTING TABLES

**Table S1** X-ray scattering factors for headgroups ($b_H^X$) and chains ($b_C^X$) of lipids used in this study, calculated as the total number of electrons of the constituent atoms and neglecting the $q$-dependence.

| Lipid | $b_H^X$ [e⁻] | $b_C^X$ [e⁻] |
|---|---|---|
| di14:1PC | 164 | 206 |
| di16:1PC | 164 | 238 |
| di18:1PC | 164 | 270 |
| di20:1PC | 164 | 302 |
| di22:1PC | 164 | 334 |

**Table S2** Structural parameters derived from analysis of experimental SAXS data. Samples are fluid-phase bilayers composed of PC lipids with two monounsaturated chains, with data collected at 25°C. $V_{HL}$ was fixed to 329 Å$^3$ (Scott 2019). Uncertainty in fitted parameters is estimated as 2% of the obtained value.

| Lipid | $V_L$ [Å$^3$] | $A_L$ [Å$^2$] | $D_B$ [Å] | $D_{HH}$ [Å] | $2D_C$ [Å] |
|---|---|---|---|---|---|
| di14:1 | 1089 ± 5 | 65.0 ± 1.3 | 33.5 ± 0.7 | 29.1 ± 0.6 | 23.4 ± 0.5 |
| di16:1 | 1195 ± 6 | 65.9 ± 1.3 | 36.2 ± 0.7 | 32.0 ± 0.6 | 26.3 ± 0.5 |
| di18:1 | 1300 ± 7 | 66.3 ± 1.3 | 39.2 ± 0.8 | 35.0 ± 0.7 | 29.3 ± 0.6 |
| di20:1 | 1406 ± 7 | 64.8 ± 1.3 | 43.4 ± 0.9 | 39.1 ± 0.8 | 33.3 ± 0.7 |
| di22:1 | 1509 ± 8 | 63.4 ± 1.3 | 47.6 ± 1.0 | 43.2 ± 0.9 | 37.2 ± 0.7 |

**Table S3** Structural parameters obtained from molecular dynamics simulations of fluid-phase bilayers. Bilayers were constructed with PC lipids containing two monounsaturated chains. Simulations were conducted at 25°C.

| Lipid | $V_L$ [Å$^3$] | $V_{HL}$ [Å$^3$] | $A_L$ [Å$^2$] | $D_B$ [Å] | $D_{HH}$ [Å] | $2D_C$ [Å] |
|---|---|---|---|---|---|---|
| di14:1 | 1061.0 | 308.8 | 68.1 | 31.2 | 28.0 | 22.3 |
| di16:1 | 1173.2 | 309.7 | 68.4 | 34.3 | 31.2 | 25.3 |
| di18:1 | 1281.5 | 309.5 | 68.0 | 37.7 | 35.2 | 28.6 |
| di20:1 | 1388.5 | 311.8 | 66.6 | 41.7 | 38.0 | 32.3 |
| di22:1 | 1496.2 | 312.7 | 64.5 | 46.4 | 42.4 | 36.7 |

**Table S4** Structural parameters derived from analysis of simulated cryoEM images of fluid-phase PC bilayers. Bilayers were composed of lipids with two monounsaturated chains. Simulations were conducted at 25°C. The true value of the trough-trough distance $D_{TT}^{true}$ is obtained by analytical projection of the electron phase shift profile of the simulated vesicle followed by smearing with a contrast transfer function as described in section S.3. The recovered trough-trough distance $D_{TT}^{rec}$ corresponds to the measured distance between the troughs of the I(w) profile obtained by averaging the profiles of 5 nm arc-length bilayer segments from noisy images of 60 simulated vesicles (i.e., the same analysis used to determine $D_{TT}$ from experimental images). Uncertainty in $D_{TT}$ obtained from image analysis was estimated using standard Monte Carlo methods (Press 2002). $D_{ves}$ is the simulated vesicle diameter.

| Lipid | $D_{ves}$ [nm] | $N_{ves}$ | $N_{seg}$ | $D_{TT}^{true}$ [Å] | $D_{TT}^{rec}$ [Å] |
|---|---|---|---|---|---|
| di14:1 | 60 | 60 | 2280 | 25.55 | 25.3 ± 0.4 |
| di16:1 | 60 | 60 | 2280 | 27.05 | 27.3 ± 0.4 |
| di18:1 | 60 | 60 | 2280 | 29.30 | 29.4 ± 0.6 |
| di20:1 | 60 | 60 | 2280 | 32.40 | 32.3 ± 0.6 |
| di22:1 | 60 | 60 | 2280 | 35.89 | 35.5 ± 0.6 |

**Table S5** Structural parameters for ternary mixtures at 25°C obtained from molecular dynamics simulations (Sodt 2014).

| Mixture | $\chi_{DPPC}$ | $\chi_{DOPC}$ | $\chi_{CHOL}$ | $V_L$ [Å$^3$] | $V_{HL}$ [Å$^3$] | $A_L$ [Å$^2$] | $D_B$ [Å] | $D_{HH}$ [Å] | $2D_C$ [Å] |
|---|---|---|---|---|---|---|---|---|---|
| Ld | 0.29 | 0.6 | 0.11 | 1165.2 | 310.4 | 57.1 | 40.8 | 40.4 | 31.2 |
| Lo | 0.55 | 0.15 | 0.30 | 991.6 | 397.3 | 40.9 | 48.5 | 46.0 | 37.9 |
